# Gadusol in epidermal mucus protects larval zebrafish from UVB-induced DNA damage

**DOI:** 10.64898/2026.10.26.762048

**Authors:** Kristin Johnson, James A. Gagnon

## Abstract

Ultraviolet B (UVB) radiation causes DNA damage and poses a threat to organisms exposed to sunlight. Zebrafish embryos are protected from UVB by maternally deposited gadusol, a small-molecule sunscreen, but it has been unclear how protection is maintained as maternal gadusol is depleted during larval development. Here, we show that gadusol synthesis in the developing zebrafish skin is required for UVB defense throughout larval development. Maternal gadusol persists through early larval stages, but juvenile fish depend solely on zygotic gadusol for protection against DNA damage and death after UVB exposure. Melanin provides a more modest contribution to survival. Gadusol is present in the epidermal mucus layer at the body’s outermost surface, positioning it to absorb UVB before it reaches underlying cells. Depletion of epidermal mucocytes increases UVB-induced DNA damage and activates a DNA damage response. Together, these findings identify a maternal-to-zygotic handoff in gadusol-based photoprotection and support a model in which gadusol in surface mucus intercepts UVB before it reaches living tissues.

## INTRODUCTION

Sunlight is an essential energy source for life on earth, providing light and heat and driving photosynthetic food webs. However, to take advantage of these energy sources, organisms are exposed to the harmful effects of ultraviolet radiation also emitted by the sun. While UVC is largely blocked by the ozone layer, UVA and UVB reach the surface of earth and can penetrate more than 10 meters into water (Van Der Leun 2004) (Karentz and Lutze 1990). UVB radiation, spanning wavelengths of light from 280 to 315 nanometers, can induce DNA and organelle damage, posing a serious risk for all living organisms. To balance the tension between access to sun-derived vital resources and damage from UVB, organisms have evolved an array of mechanisms to correct or avoid UVB-induced DNA damage. Cells can detect and repair DNA damage after UVB, but all repair mechanisms are imperfect (Chatterjee and Walker 2017).

Sunscreens prevent DNA damage by reflecting, scattering, or absorbing UV photons (Cockell and Knowland 1999). A diverse molecular arsenal of sunscreens are produced by living organisms (Gao and Garcia-Pichel 2011). Melanin, a pigment produced by skin melanocytes that absorbs and scatters UVB, is the best known vertebrate sunscreen (Brenner and Hearing 2008). However, an alternative sunscreening molecule is produced by many other vertebrates. Gadusol, a small molecule that can absorb UVB, was originally discovered in cod eggs (Plack et al. 1981). Gadusol has a molecular structure similar to but distinct from mycosporine-like amino acids (MAAs), a class of sunscreen compounds found in algae (Görünmek et al. 2024), microbes (Gao and Garcia-Pichel 2011), and coral (Dunlap and Shick 1998). Historically, the presence of gadusol in vertebrates was ascribed to accumulation from dietary sources or symbionts (Gao and Garcia-Pichel 2011).

Decades later, genes encoding a biosynthetic pathway for gadusol production were discovered in many vertebrate genomes (Osborn et al. 2015). Two enzymes, EEVS and MT-Ox, convert sedoheptulose 7-phosphate, a product of the pentose phosphate pathway, into gadusol (Fig. 1A). In zebrafish, gadusol is synthesized during oogenesis and maternally deposited into zebrafish eggs (Rice et al. 2023). Using *eevs* mutants that disrupt gadusol synthesis, we previously found that maternally-provided gadusol protects zebrafish embryos and larvae from UVB during the first five days of development. In the absence of gadusol, mutant embryos accumulate DNA damage, have an elevated DNA damage response and increased apoptosis, and ultimately do not survive a physiologically relevant UVB challenge.

**Figure 1.**
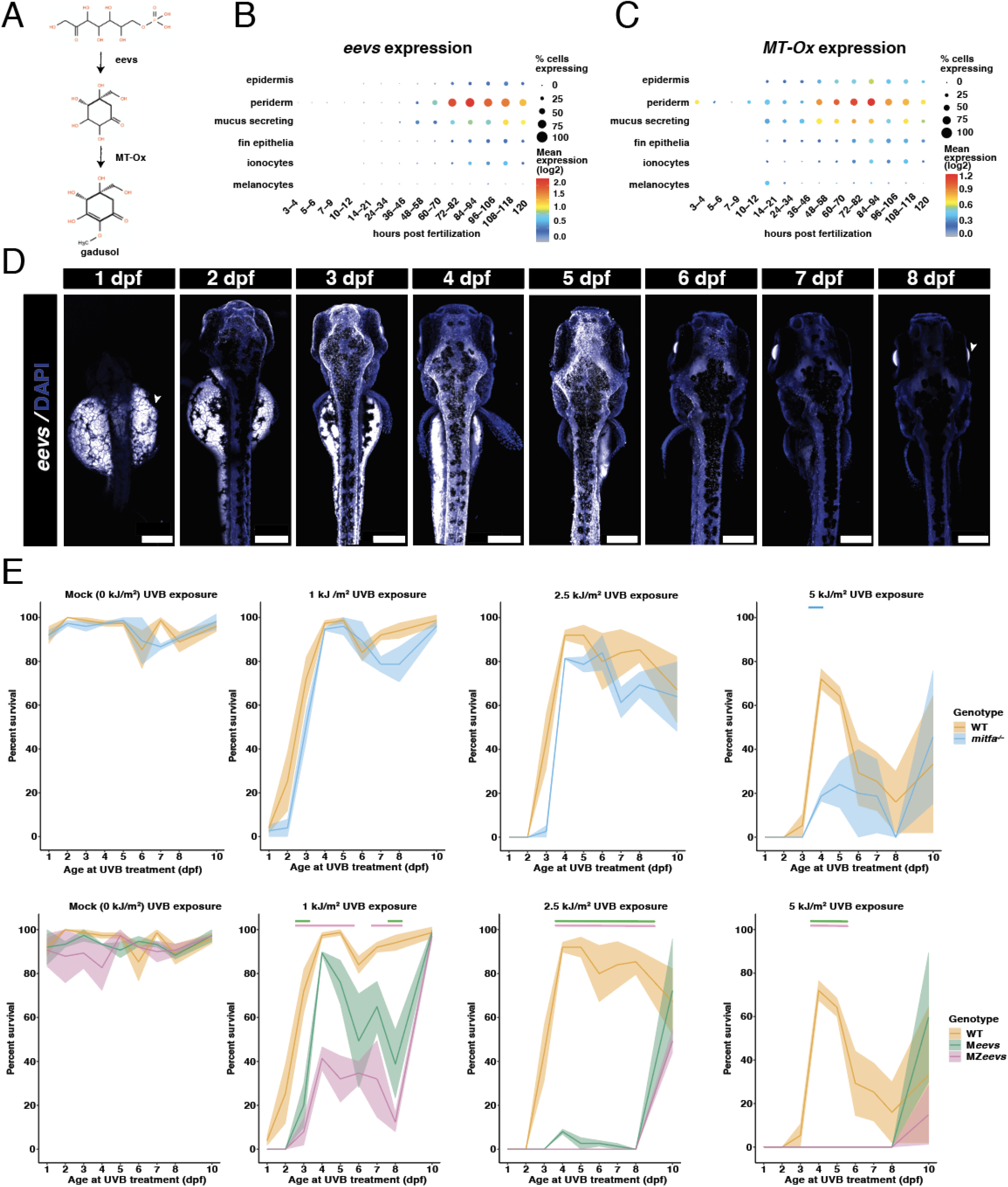
*Z*ygotic gadusol production provides protection against UVB in zebrafish larvae. (A) The gadusol biosynthesis pathway. (B) Single-cell expression of *eevs* in skin cell types through the first five days of development, visualized with DaniocellDesktop (Evans and Farrell 2025). (C) Single-cell expression of *MT-Ox* in skin cell types through the first five days of development, visualized as above. (D) Representative images of HCR-Pro RNA FISH for *eevs* (white) counterstained with DAPI (blue) through the first 8 days of development. White bars are 200 uM scale bars. White arrows indicate autofluorescence captured in the yolk and eye lens. (E) Survival after a UVB challenge through zebrafish development. Each time point on the x-axis represents the day of development when fish were treated with 0 kJ/m^2^, 1 kJ/m^2^, 2.5 kJ/m^2^, or 5 kJ/m^2^ UVB. Fish survival was scored at 21 dpf. Shading represents SEM. n= 75, C=3 for each genotype at each time point. Bars at the top of plots represent a statistically significant difference (p < 0.05), between each genotype and WT (blue for *mitfa*^-/-^, green for M*eevs* and pink for MZ*eevs*), calculated using a three way ANOVA followed by Tukey’s HSD. Full list of p-values is provided as Supplemental Table S1.

Some evidence suggests that sunscreens may also protect later stages of zebrafish development. Larval melanophores respond to UVB by increasing whole-body pigmentation (Mueller and Neuhauss 2014), and internal melanocytes positioned over the larval zebrafish kidney can protect HSPCs from UV-induced DNA damage (Kapp et al. 2018). However, melanophores do not cover the entire surface of the larval zebrafish, and larvae lacking melanophores have only a modest decrease in survival upon UVB treatment (Rice et al. 2023). The *eevs* and *MT-Ox* genes are transcribed during the first days of embryo development (Osborn et al. 2015), suggesting that gadusol synthesis may initiate at the embryo-to-larval transition to provide an alternative sunscreen to melanin.

The best place for a sunscreen is the external surface of the body, where it could block living cells from UVB. Teleost fish epidermis is the primary physiological barrier to external hazards (Hawkes 1974). Epidermal goblet cells, referred to here as mucocytes, release mucins, antimicrobial molecules, and other factors, creating a dynamic external layer of mucus that is the superficial surface of the body in direct contact with the environment. This mucus layer is formed by 4 days post-fertilization in zebrafish larvae (Lu et al. 2021). MAAs have been found in the mucus of both fish (Braun et al. 2016) and coral (Teai et al. 1998), and mucosal MAA concentration has been correlated with UVB protection (Braun et al. 2016). It has also been shown that some fish species increase mucus production in response to UVB exposure (Zamzow et al. 2013). Together, these studies suggest that fish mucus may serve a role as a sunscreen.

Here, we evaluated the hypothesis that gadusol is produced in the zebrafish skin and secreted into mucus to protect larvae from UVB-induced DNA damage. We found that *eevs* and *MT-Ox* are transcribed in larval skin cells, including epidermal mucocytes. Gadusol produced in mucocytes is present in mucus and is required to prevent DNA damage and death in zebrafish larvae and juveniles. Zebrafish depleted of mucocytes lack superficial gadusol and are sensitized to UVB-induced DNA damage. Melanin serves a secondary role as a sunscreen at these stages. Together, these data demonstrate that zebrafish larvae protect themselves from UVB with gadusol in their surface mucus as a sunscreen before it can reach living tissues.

## RESULTS

### The gadusol synthesis enzyme *eevs* is expressed in the skin

Gadusol synthesis genes have previously been reported to be expressed in the zebrafish ovary and during embryo development, and gadusol synthesized de novo begins to accumulate in larvae (Osborn et al. 2015; Rice et al. 2023). To determine where gadusol might be synthesized in the larval zebrafish, we analyzed published single-cell RNA sequencing (scRNAseq) data through the first five days of development (Sur et al. 2023). We found that the gadusol synthesis enzyme *eevs* is transcribed zygotically starting at 2 days post-fertilization (dpf) through 5 dpf in the periderm, epidermis, fin epithelia, ionocytes and mucus-secreting cells, with little to no expression outside of these cell types (Fig. 1B, S1A). *MT-Ox* is also expressed in skin and mucus-secreting cells in early development, alongside broader expression outside the skin (Fig. 1C, S1B). Consistent with these data, examination of another zebrafish scRNAseq atlas also revealed expression of both *eevs* and *MT-Ox* in the periderm at 3 dpf and 5 dpf (Fig. S1C) (Lange et al. 2024). To confirm the *eevs* expression pattern in the larval zebrafish, we used hybridization chain reaction fluorescent in situ hybridization (HCR-FISH) to visualize *eevs* mRNA across a time course of zebrafish development. Consistent with the scRNAseq data, *eevs* was expressed throughout the skin starting at 2 dpf and peaking between 3 and 5 dpf (Fig. 1D, S1D). Together, these data show that gadusol synthesis genes are expressed throughout the larval zebrafish skin.

We hypothesized that gadusol synthesis in the larval zebrafish skin would provide protection from UVB. To assess UVB protection, we systematically analyzed the survival of larval zebrafish of various genotypes after doses of UVB irradiation delivered at different developmental ages. The genotypes we compared were wild-type (WT), *mitfa*^*-/-*^ (fish lacking melanophores, which produce all skin melanin), MZ*eevs* (*eevs*^*-/-*^ fish lacking both maternally provided and zygotically produced gadusol), and M*eevs* (*eevs*^*+/-*^, fish lacking maternally provided gadusol but capable of zygotic gadusol production). We treated all fish with a single dose of UVB at four dosages (0 kJ/m^2^ (Mock), 1 kJ/m^2^, 2.5 kJ/m^2^, 5 kJ/m^2^) at days 1, 2, 3, 4, 5, 6, 7, 8 or 10 of larval development. Each condition was evaluated in triplicate. For each group, we scored endpoint survivorship at 21 dpf, a time at which fish must be capable of swimming and feeding independently to survive. This experiment scored >2,500 fish across all treatment groups.

As expected, there was no difference in survival between genotypes after mock UVB exposure at any developmental time point (0 kJ/m^2^, Fig. 1E, left column). Across all genotypes, increased UVB exposure led to decreased survival, with substantial variation depending on the developmental time point of UVB exposure, as expected. Embryos at 1, 2 or 3 dpf were most sensitive to UVB exposure, and resilience increased dramatically starting at 4 dpf. All genotypes showed more tolerance for UVB at 4-5 dpf compared to 7-8 dpf, consistent with a previous study (Keating et al. 2026).

MZ*eevs* animals were more sensitive to UVB compared to animals of all other genotypes after 1 kJ/m^2^ UVB exposure at 4-8 dpf (Fig. 1E, left-center column). This matched the days following peak zygotic *eevs* expression (Fig. 1B,D). Fish treated with 2.5 kJ/m^2^ UVB had a significant survival difference between those with maternally provided gadusol (WT) and without maternally provided gadusol (MZ*eevs* and M*eevs*) (Fig. 1E, right-center column), suggesting that maternally provided gadusol still participates in UVB protection through 8 dpf. Across all timepoints and UVB doses, fish lacking melanin (mitfa^-/-^) had little or no survival difference when compared to WT, with the exception of 5 kJ/m^2^ at 4 dpf (Fig. 1E, top row), suggesting a secondary role for melanin as a sunscreen at these stages. Together, these data suggest that zygotically produced gadusol has an important role in protecting fish from UVB as early as 4 dpf, even as maternally provided gadusol continues to participate in UVB defense (Rice et al. 2023).

### The maternal-to-zygotic handoff of gadusol production occurs by 21 dpf

Our data indicate gadusol is a remarkably stable molecule - maternally provided gadusol persists and participates in UVB defense for nearly a week after fertilization. At the same time, zygotic production of gadusol commences and begins contributing to UVB defense. To determine when the handoff between maternally provided and zygotically produced gadusol is complete, we dosed later stages of zebrafish larvae with UVB. First, we treated 8 dpf WT, MZ*eevs* and M*eevs* fish with 1 kJ/m^2^ UVB and scored their survival at 21 dpf (Fig. 2A). Consistent with our previous experiment, less than 10 percent of MZ*eevs* fish survived a 1 kJ/m^2^ UVB exposure, while over 40% of M*eevs* survived and over 90% of WT fish survived (Fig. 2B). These data confirm that both maternally provided and zygotically produced gadusol work together as sunscreens to protect this developmental timepoint. Next, we treated 21 dpf WT, M*eevs* and MZ*eevs* fish with 10 kJ/m^2^ of UVB and scored their survival at 23 dpf (Fig. 2A). At this time point, M*eevs* have no survival deficit compared to WT, suggesting that by 21 dpf maternally provided gadusol plays no role in UVB defense, perhaps due to depletion. However, only ∼50% of MZ*eevs* fish survived this dose of UVB, significantly less than both WT (p = 0.0025) and M*eevs* (p = 0.0094), suggesting that juvenile fish still depend on zygotically produced gadusol for UVB defense (Fig. 2C). Finally, we asked whether melanin participates in UVB defense in juvenile fish. By 21 dpf, zebrafish have developed markedly more melanin and formed the characteristic stripes of the lateral body that give zebrafish their name. We treated 21 dpf WT and *mitfa*^*-/-*^ fish, which lack melanophores, with 10 kJ/m^2^ UVB and scored their survival at 23 dpf. There was a modest but significant reduction in UVB tolerance between WT and *mitfa*^*-/-*^ fish (p = 0.0493) - ∼75% of fish that lack melanin survived this UVB dose compared to nearly 100% of the WT, suggesting melanin acts as a secondary sunscreen at this time point (Fig. 2D).

**Figure 2.**
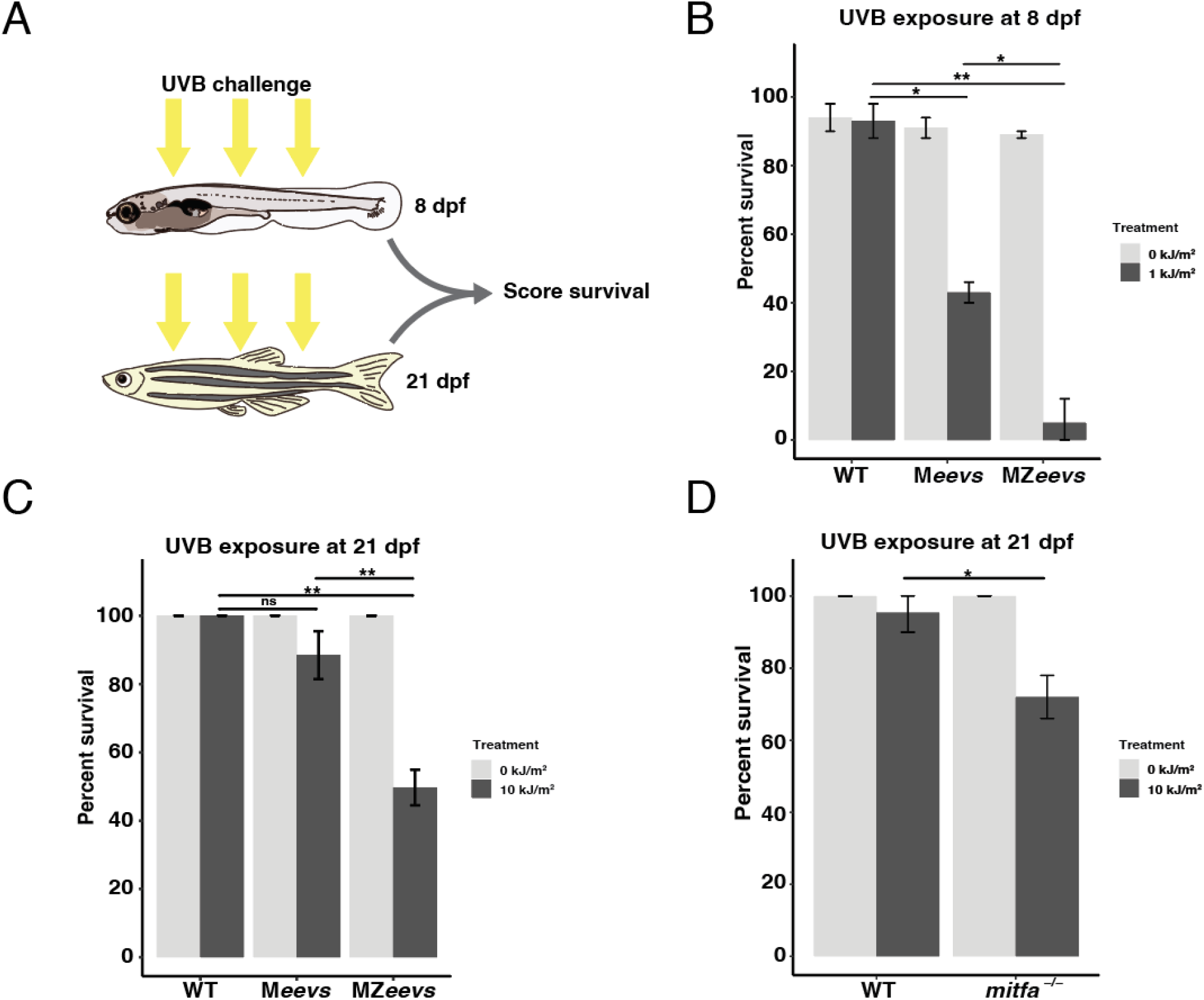
Developing zebrafish rely on both maternally provided and zygotically produced gadusol. (A) Schematic of UVB challenge experiments. (B) Survival scored at 21 dpf after 8 dpf WT, M*eevs* and MZ*eevs* larval zebrafish were treated with 1 kJ/m^2^ UVB. Error bars represent SEM, significance calculated using Student’s t-test, *WT - M*eevs* p = 0.0194, *M*eevs* - MZ*eevs* p = 0.0379 **WT - MZ*eevs* p = 0.0019, n=100, C=2 for each genotype. (C) Survival scored at 23 dpf after 21 dpf WT, M*eevs* and MZ*eevs* larval zebrafish were treated with 10 kJ/m^2^ UVB. Error bars represent SEM, statistical significance calculated using Student’s t-test, ns WT-M*eevs* p = .1518, ** WT - MZ*eevs* p = 0.0025, **M*eevs* – Mz*eevs* p = 0.0094, n=97, 95, 98, 95, 94, 95, C=2. (D) Survival scored at 23 dpf after 21 dpf WT and *mitfa*^-/-^ larval zebrafish were treated with 10 kJ/m^2^ UVB. Error bars represent SEM, significance calculated using Student’s t-test * p = 0.0493, n=95, 91, 92, 91 C=2.

### Zygotically produced gadusol acts as a sunscreen by preventing DNA damage

We previously found that gadusol acts as a sunscreen in 1 dpf embryos by preventing UVB-induced DNA damage, notably cyclobutane pyrimidine dimers (CPDs), the specific type of DNA damage induced by UVB (Rice et al. 2023). To determine whether zygotically produced gadusol in the skin also prevents DNA damage, we used an antibody specific for CPDs (TDM-2 antibody) (Rice et al. 2023). We treated 8 dpf WT, *mitfa*^*-/-*^, M*eevs* and MZ*eevs* larvae with 1 kJ/m^2^ UVB and collected immediately after treatment, so as not to allow time for DNA damage repair, then proceeded with immunostaining with the CPD antibody. MZ*eevs* fish accumulated more DNA damage than WT (p = 0.00007), *mitfa*^*-/-*^ (p = 0.00002) or M*eevs* (p = 0.0115) fish, consistent with the hypothesis that zygotic production of gadusol is required for UVB defense (Fig. 3A, B). Imaging UVB-treated larvae from dorsal and lateral views revealed that MZ*eevs* fish accumulated DNA damage on the dorsal side of the fish, particularly the head, eye, fins and lateral line with minimal DNA damage on the ventral side of the body (Fig 3C). This is consistent with the UVB illumination coming from above the fish, similar to exposure in the wild. WT fish show far less DNA damage, mainly in the tail fin and above the swim bladder, consistent with gadusol effectively absorbing UVB on the dorsal surface of WT fish.

**Figure 3.**
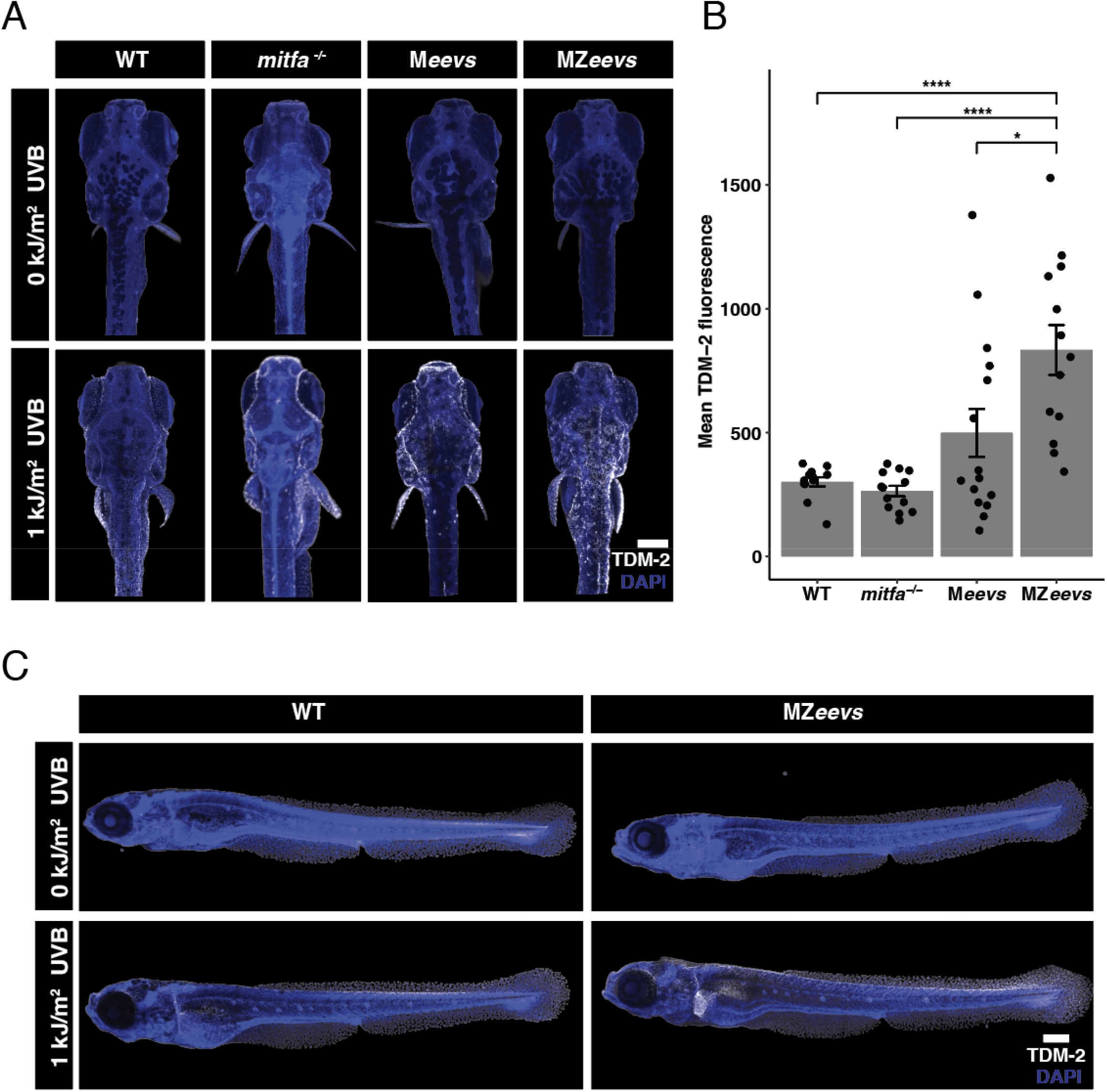
Zygotically produced gadusol prevents DNA damage. (A) Representative dorsal view images of immunohistochemistry for TDM-2 (white), an antibody that recognizes CPDs, counterstained with DAPI (blue). 8 dpf larval zebrafish were exposed to 1 kJ/m^2^ UVB, or mock exposure, fixed immediately and stained. White bar is a 200 uM scale bar for all images. (B) Quantification of mean TDM-2 fluorescence, with background subtracted, relative to UV negative control. Error bars are SEM, significance calculated using one way ANOVA followed by Tukey’s HSD, WT - MZ*eevs* ****p = 0.00007, *mitfa*^*-/-*^ - MZ*eevs* ****p = 0.00002, M*eevs* - MZ*eevs* *p = 0.0115, n = 12, 13, 15, 13, C = 5 for all. (C) Representative lateral view images of immunohistochemistry for TDM-2 (white), counterstained with DAPI (blue). 8 dpf larval zebrafish were exposed to 1 kJ/m^2^ UVB, or mock UV exposure, fixed immediately and stained. White bar is a 200 uM scale bar for all images.

### Gadusol is present in epidermal mucus to prevent DNA damage

Based on the expression of *eevs* and *MT-Ox*, we hypothesized that gadusol is being produced in skin cells, particularly the epidermal mucocytes that produce superficial mucus as an environmental barrier (Lu et al. 2021). Since there is no available antibody or other staining reagent to mark gadusol, we used a previously-established indirect measure of gadusol abundance - UVB absorbance (Osborn et al. 2015) (Rice et al. 2023). To determine if gadusol is present in epidermal mucus, we collected mucus by forceps scraping and indirectly measured gadusol absorbance using a spectrophotometer to measure UVB absorbance (Fig. 4A, top). Using this assay, we infer that gadusol is present in mucus of WT fish but not MZ*eevs* fish (p = 5.607E-06) (Fig. 4A, bottom).

**Figure 4.**
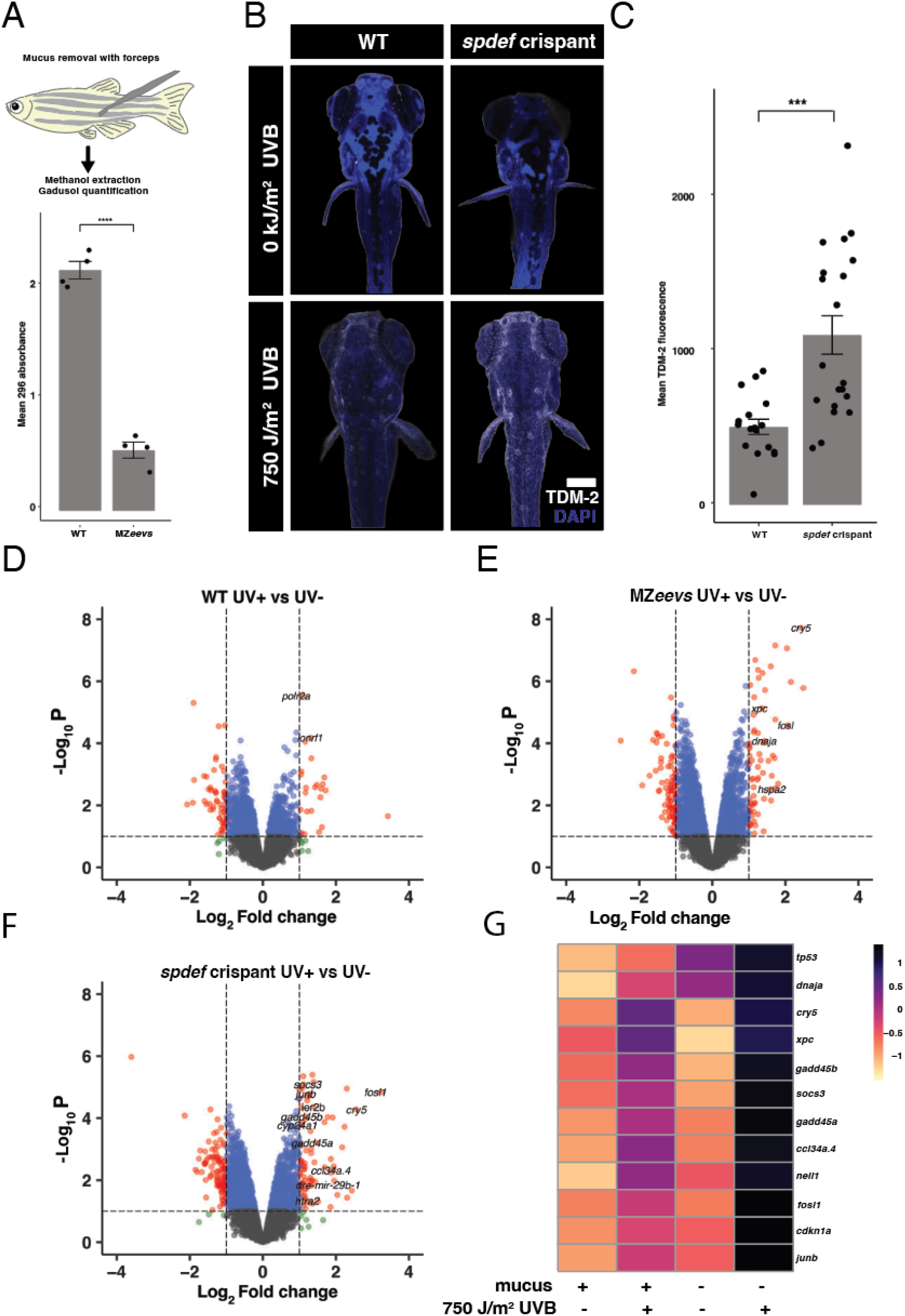
Gadusol in mucus protects fish from UVB. (A, top) Mucus was removed from the lateral body surface by scraping with forceps and extracted with methanol. Quantification of gadusol in mucus extracts, using absorbance at 296 nm normalized to 260 nm absorbance (DNA). Error bars are SEM, significance calculated using student’s t-test **** p = 5.607E-0,. (B) Representative images of immunohistochemistry for TDM-2 (white), an antibody that recognizes CPDs counterstained with DAPI (blue). 6 dpf WT or *spdef* crispant larval zebrafish were exposed to 750 J/m^2^ UVB, or mock UV exposure, fixed immediately and stained. White bar is a 200 uM scale bar for all images. (C) Quantification of mean TDM2 fluorescence, with background subtracted, relative to no UV control for each condition. n = 17, 20 C = 3 for all. Error bars are SEM, significance calculated using student’s t-test ***p = 0.0002. (D-F) Comparisons of RNAseq data represented using volcano plots showing differentially expressed genes in UVB treated vs untreated samples in 6 dpf WT, MZ*eevs* and *spdef* crispant fish. Significantly differentially expressed genes are in red, select DNA damage response genes are named. RNA was collected 5 hours post mock or 750 J/m^2^ UVB exposure. (G) Significant upregulation of select UVB response and DNA damage GO term-associated genes measured from the indicated conditions and genotypes. Gene expression is scaled by rows.

We hypothesized that gadusol in epidermal mucus was providing a sunscreen located outside living tissues. To manipulate mucus production, we identified *spdef* as a transcription factor specifically expressed in zebrafish mucus-secreting cells at early developmental stages (Fig. S2A). Spdef is known to regulate epithelial mucocyte differentiation in mice (Chen et al. 2009; Rajavelu et al. 2015) and is highly coexpressed in epidermal mucocytes with the epidermal specific mucin genes *muc5*.*1* and *muc5*.*2* (Fig. S2B). We used CRISPR-Cas9 to generate crispants for *spdef* by targeting guide RNAs to the functional domains of exon 2 and exon 4 (Fig. S2C). Alcian blue staining revealed a significant reduction in epidermal mucocytes in *spdef* crispants as indicated by lack of binding to acidic mucin molecules (Fig. S2D,E) (Chuang et al. 2019). RNAseq of 6 dpf WT and *spdef* crispant fish suggests a strong depletion of mucus proteins as indicated by the near-total loss of mucins expression in *spdef* crispants compared to control fish, including *muc5*.*1 and muc5*.*2* which are specific to epidermal mucus (Jevtov et al. 2014) (Fig. S2F). To determine that mucus is indeed necessary to prevent UVB-induced DNA damage, we exposed 6 dpf WT and *spdef* crispants to 750 J/m^2^ UVB and immediately collected them for immunofluorescence for CPDs, as before. *spdef* crispant fish lacking mucus had elevated levels of DNA damage compared to WT fish (p = 0.0002) (Fig. 4B,C). We characterized the transcriptional response to UVB in the absence of mucus by comparing RNAseq of *spdef* crispants and WT larvae after UVB exposure. Five hours after UVB exposure, we observed a modest upregulation of DNA damage response genes in WT fish exposed to UVB compared to unexposed controls (Fig. 4D). A much greater number of genes was upregulated in both fish lacking gadusol (*MZeevs*) (Fig. 4E) and fish lacking mucus (*spdef* crispants) (Fig. 4F), both of which were strongly enriched for DNA damage response and repair genes. Further analysis revealed that *spdef* crispants had significantly higher expression of many key stress response and DNA repair genes (e.g., *tp53, gadd45a, cry5, ddb2*) relative to UVB-treated WT larvae (Fig. 4G). Together, these data suggest that gadusol is present in epidermal mucus to act as a sunscreen and protect fish from UVB-induced DNA damage and cellular stress.

## DISCUSSION

Animals must protect themselves from UVB damage throughout their lives. We previously found that maternally provided gadusol serves as the primary sunscreen in zebrafish through the first five days of development (Rice et al. 2023). However, this maternal supply is inevitably depleted with time. Here, we show that the gadusol synthesis pathway is activated in the larval zebrafish skin. This zygotically produced gadusol contributes to sun protection starting at day 4 of development, initially supplementing the protection provided by maternal gadusol, and continuing until at least 21 days of age, when it is the primary sunscreen. This zygotically produced gadusol is present in epidermal mucus, providing sun protection at the external surface of the body. CRISPR-mediated disruption of mucocyte differentiation, which blocks epidermal mucus production, made larval fish more sensitive to UVB-induced DNA damage. Together, these data show that gadusol in epidermal mucus is necessary for prevention of UVB-induced DNA damage in developing zebrafish.

Gadusol is a remarkably stable molecule. Maternally provided gadusol persists as a sunscreen over a week into development. At the same time, zygotic gadusol production begins, providing overlapping protection for larval and juvenile stages. The stability of gadusol may partially mitigate the metabolic cost of its synthesis if gadusol only needs to be produced periodically to maintain enough gadusol to effectively protect against UVB. Others have estimated that microbes may allocate >10% of their metabolic activity towards sunscreen protection (Gao and Garcia-Pichel 2011). The repeated loss of gadusol synthesis enzymes in teleost species not exposed to the sun (Rice et al. 2023) also suggest an evolutionary tradeoff to gadusol production. Gadusol is a transparent small molecule that cannot be visualized directly - no antibody or other reagent for labeling is available. It is possible that heavy isotope labeling of a gadusol precursor through feeding and LC-MS could permit studies of gadusol persistence, stability, and response to UVB.

Zygotically produced gadusol is an important sunscreen in juvenile zebrafish. However, juvenile animals also increase their tolerance to UVB independent of gadusol. UVB penetrates only superficial tissues, so the growth of developing fish inevitably means that UVB can only induce damage in a proportionally smaller fraction of overall cells and thus animals become more tolerant to UVB as they grow. Additionally, we found that melanin is a secondary sunscreen at this stage, but many other untested factors may contribute to UVB tolerance.

Other pigment cells in zebrafish such as iridophores and xanthophores can reflect and absorb light, yet their potential roles in UVB defense have not been investigated. Finally, physical structures (e.g., scales) and behavioral traits (e.g., preference for deeper waters) may play important developmental roles in UVB defense. A holistic survey of many physiologically-relevant defense strategies against UVB may be important as climate change drives fish into new habitats on Earth.

We do not know whether UVB exposure may cause increased gadusol synthesis or mucus deposition. It has long been known that UVB exposure increases melanocyte production in human skin (Stierner et al. 1989), and more recently drives melanophore expansion in larval zebrafish (Mueller and Neuhauss 2014). It will be interesting to see if fish exhibit a similar response with an upregulation of *eevs* and increase in gadusol synthesis in response to UVB. Alternatively, zebrafish may respond to UVB exposure with an increase in secretion of gadusol into the mucus, similar to what is seen with MAAs in some species of reef fish (Braun et al. 2016), (Zamzow et al. 2013). It will be interesting to determine if zebrafish increase production or secretion of gadusol in response to UVB, and if such an increase protects fish from future UVB exposures.

Gadusol synthesis genes are found in the genomes of fish, birds, reptiles, and amphibians (Osborn et al. 2015). However, fish are the only class of vertebrates where gadusol has been directly detected. There have been many independent losses of gadusol in fish species not exposed to the sun, suggesting that the primary role of gadusol is photoprotection (Rice et al. 2023). While no functions have been ascribed to gadusol besides UVB defense, *eevs* is expressed in other organs, including the liver (Aniseia et al. 2026). It remains possible that gadusol may have other roles, similar to how melanin is involved in signaling, thermoregulation and immunity (McNamara et al. 2021). It will be fascinating to see whether gadusol is employed for UVB defense or other functions in non-fish vertebrates, and how gadusol production may connect to the transition from aquatic to terrestrial life hundreds of millions of years ago.

## METHODS

### Zebrafish husbandry

All zebrafish lines were maintained in accordance with standard husbandry practices at the CBRZ zebrafish facility at the University of Utah. This study was conducted with approval of the Office of Institutional Animal Care and Use Committee of the University of Utah’s animal care and use program.

### Gadusol extractions and detection via nanodrop

To confirm the presence or absence of gadusol for each genotype in each experiment, we measured absorbance at 296 nm of polar compounds extracted from pooled embryos using Nanodrop UV-vis spectrophotometry as previously described (Rice et al. 2023). Briefly, 25 embryos/larvae were placed in a microfuge tube. All excess water was removed with a Pasteur pipette. 100 ml of 80:20 (v:v) methanol:water was added to embryos. Embryos were mashed with a microfuge pestle for 15 s. Samples were left to extract for at least 15 min, and then centrifuged at 12,000 g. Clear supernatant, containing polar compounds such as gadusol, was separated for analysis by spectrophotometry. Epidermal mucus was removed by scraping the lateral surface of 6 adult fish with sterile forceps and placing mucus into 50 uL of 80% EtOH. Polar molecules were extracted and analyzed as described above. Gadusol was quantified by measuring absorbance at 296 nm and normalizing to absorbance at 260 nm.

### UV exposure and survival assays

A raised and inverted UVP transilluminator with 306 nm broadband UVB bulbs was used (Ushio G8T5E) on the ‘‘low’’ setting, as previously described (Rice et al. 2023). UVB doses were calibrated using a radiometer (Solarmeter UVB) at a fluence rate of 2.5 W/m^2^ in 30 mL of clear E3 media. For time course experiments, groups of 25 fish were exposed to 0 kJ/m^2^ (mock exposure), 1 kJ/m^2^, 2.5 kJ/m^2^, or 5 kJ/m^2^ at 1, 2, 3, 4, 5, 8 and 10 dpf. All survival percentage data and p-values are included in Supplemental Table S1. For 8 dpf WT, M*eevs*, MZ*eevs* survival experiments in Figure 2, 8 dpf larvae were exposed to 0 kJ/m^2^ or 1 kJ/m^2^ of UVB. This is a conservative estimate of a physiologically relevant UVB dose that fish embryos would routinely experience in the wild (Behrendt et al. 2010). Survival for all of these experiments was scored at 21 days post-fertilization to ensure that all living juveniles could feed on their own. For 21 dpf survival experiments in Figure 2, 21 dpf fish were exposed to 10 kJ/m^2^ UVB and survival was scored two days later at 23 dpf. For WT versus *spdef* crispant experiments, 6 dpf fish were exposed to 750 J/m^2^ UVB and then fixed immediately for IF or collected 5 hours post-UVB exposure for RNA extraction and RNASeq.

### HCR-FISH

Larvae were collected at the designated stage and fixed in 4% PFA overnight at 4° C. Fixed larvae were rinsed in PBS and dehydrated in methanol and stored at -20° C until HCR™ was performed on all samples from a single experiment simultaneously. HCR™ (Molecular Instruments) was performed using the protocol for HCR™ Pro RNA-FISH Protocol: Sample in Solution for whole-mount embryos. Tyramide Conjugate Alexa Fluor 546 was used for fluorescent reporter detection. The *Danio rerio eevs* probe was custom designed by Molecular Instruments. Samples were mounted in 1.3% low melt agarose and imaged on a Zeiss 880 inverted confocal microscope with a 10X objective. Although HCR is widely used for RNA-FISH, the specificity of the probes used here was not independently validated.

### IF imaging and quantification of CPDs in 6dpf and 8dpf larvae

Larvae were exposed to 1 kJ/m^2^ of UVB (8 dpf) or 750 J/m^2^ UVB (6 dpf) and then immediately fixed after exposure in 4% PFA for 1 hour at 25° C. Embryos were kept in the dark during fixation to avoid photoreactivation. Fixed embryos were then washed in PBST, dehydrated in methanol, and stored at -20° C. Embryos were rehydrated into PBST and exposed to 2M HCl for 1 hour to break apart dsDNA and expose CPD epitopes, as previously described (Rice et al. 2023). Samples were blocked in 10% NGS + 5% BSA in PBST. Mouse anti-CPD primary antibody (TDM-2, Cosmo Bio) was used to stain for CPDs at 1:500 overnight at 4° C. Goat anti-mouse AF546 secondary antibody (Invitrogen) was used at 1:250 for 2 hours at 25° C to visualize CPDs. Embryos were also stained with DAPI overnight in PBS at 4° C to visualize nuclei. Larvae were mounted in 1.3% low melt agarose and imaged on a Zeiss 880 inverted confocal microscope with a 10X objective. Images were analyzed using ImageJ (Schindelin et al. 2012). To determine mean fluorescence intensity, mean pixel intensity was calculated using the sum of z-stacks over the sample area, defined by tracing around the outline of the sample, background fluorescence was subtracted from each image and mean fluorescence of UV-treated conditions was normalized to the average mean fluorescence of UV negative controls for each genotype.

### Generation of *spdef* crispants

*spdef* crispants were generated using CRISPR-Cas9 mutagenesis. Four gRNAs were designed using CRISPOR (Concordet and Haeussler 2018), two guides targeting the functional domain in exon 2 and two guides targeting the functional domain in exon 4. Guide RNAs were synthesized from DNA oligos using standard protocols (Takasugi et al. 2022). Freshly laid wild-type TU-strain embryos were injected with SpCas9 protein (NEB) mixed with gRNAs (∼300 ng/ml), KCl, and phenol red. 1-2 nL were injected into each embryo. WT controls were injected with only spCas9 protein.

### Alcian blue staining

Larvae were fixed in 4% PFA for 2 hours at 25° C, rinsed 3 x 5 min in PBS, rinsed 3 x 5 min in EtOH, dehydrated in MeOH, and stored at -20°C until processing. Larvae were rehydrated and rinsed in molecular grade water and stained in 0.01% alcian blue in 3% acetic acid for 2 hours at 25° C, rinsed in molecular grade water and rinsed in 70% EtOH 1% HCl before scoring and imaging.

### RNASeq sample preparation and analysis

Pools of 20 larvae were collected in triplicate 5 hours after mock or 750 J/m^2^ UVB exposure and homogenized with a microfuge pestle (MTC Bio). RNA was extracted using TRI Reagent (Zymo) and purified via Direct-zol RNA Miniprep Plus (Zymo). Samples were submitted to Plasmidsaurus RNASeq, 3’ end counting bulk RNA-Seq from purified RNA. Purified RNA samples were submitted to Plasmidsaurus in Seqguard Dino Preserve™ and 10X TE buffer. Count and cpm data and DESeq genes were generated by Plasmidsaurus. Volcano plots were generated with Enhanced Volcano (Blighe et al. 2019) in RStudio and heatmaps were generated with ggplot2 (Wickham et al. 2019) in RStudio.

## ACKNOWLEDGEMENTS

We thank all members of the Gagnon lab for discussions and comments. We thank Julie Hollien for help with equipment. We thank the Centralized Zebrafish Animal Resource (CZAR) and CBRZ staff for zebrafish care. We thank the Cell Imaging core at the University of Utah, specifically Xiang Wang. This research was conducted on the traditional and ancestral homeland of the Shoshone, Paiute, Goshute, and Ute Tribes. We affirm and support the University of Utah’s partnership with Native Nations and Urban Indian communities. This project was supported by National Institutes of Health grants T32HD007491 (K.J.), F32GM154475 (K.J.), R35GM142950 (J.A.G.), and by startup funds from the University of Utah Henry Eyring Center for Cell and Genome Science (J.A.G.).

---

**Figure S1.**
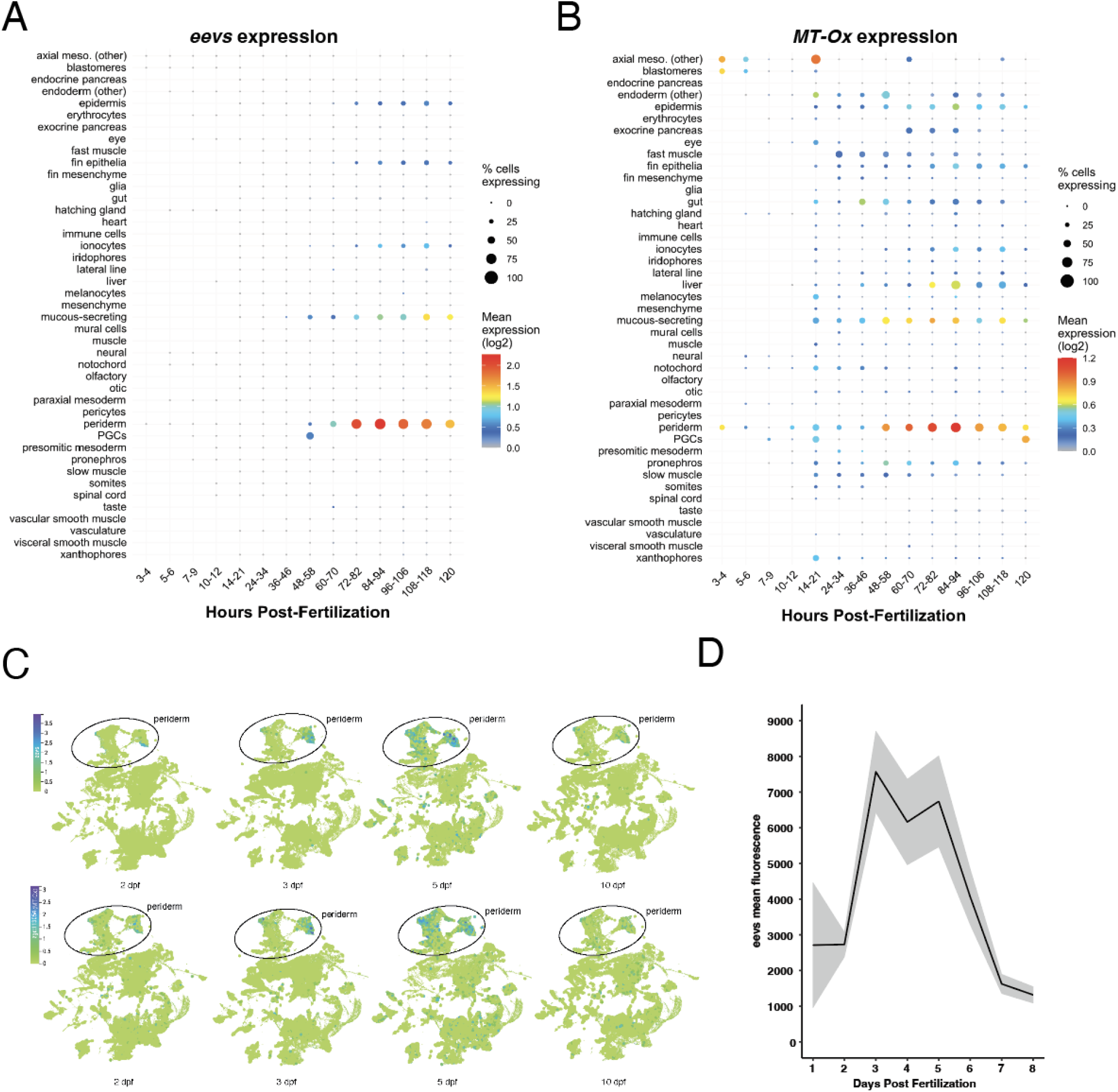
*Expression of eevs* and *MT-Ox* in the skin in early development. (A) Expression of *eevs* throughout the first five days of development. Adapted from Daniocell (Sur et al. 2023). (B) Expression of *MT-Ox* throughout the first five days of development. Adapted from Daniocell (Sur et al. 2023). (C) *Eevs* and *MT-Ox* expression data in early development shows a peak in expression in the periderm at 3-5 dpf. Adapted from Zebrahub (Lange et al 2024). (D) Quantification of mean total fluorescence of *eevs* expression with background subtracted. Error bars represent SEM, n = 3 at each timepoint.

**Figure S2.**
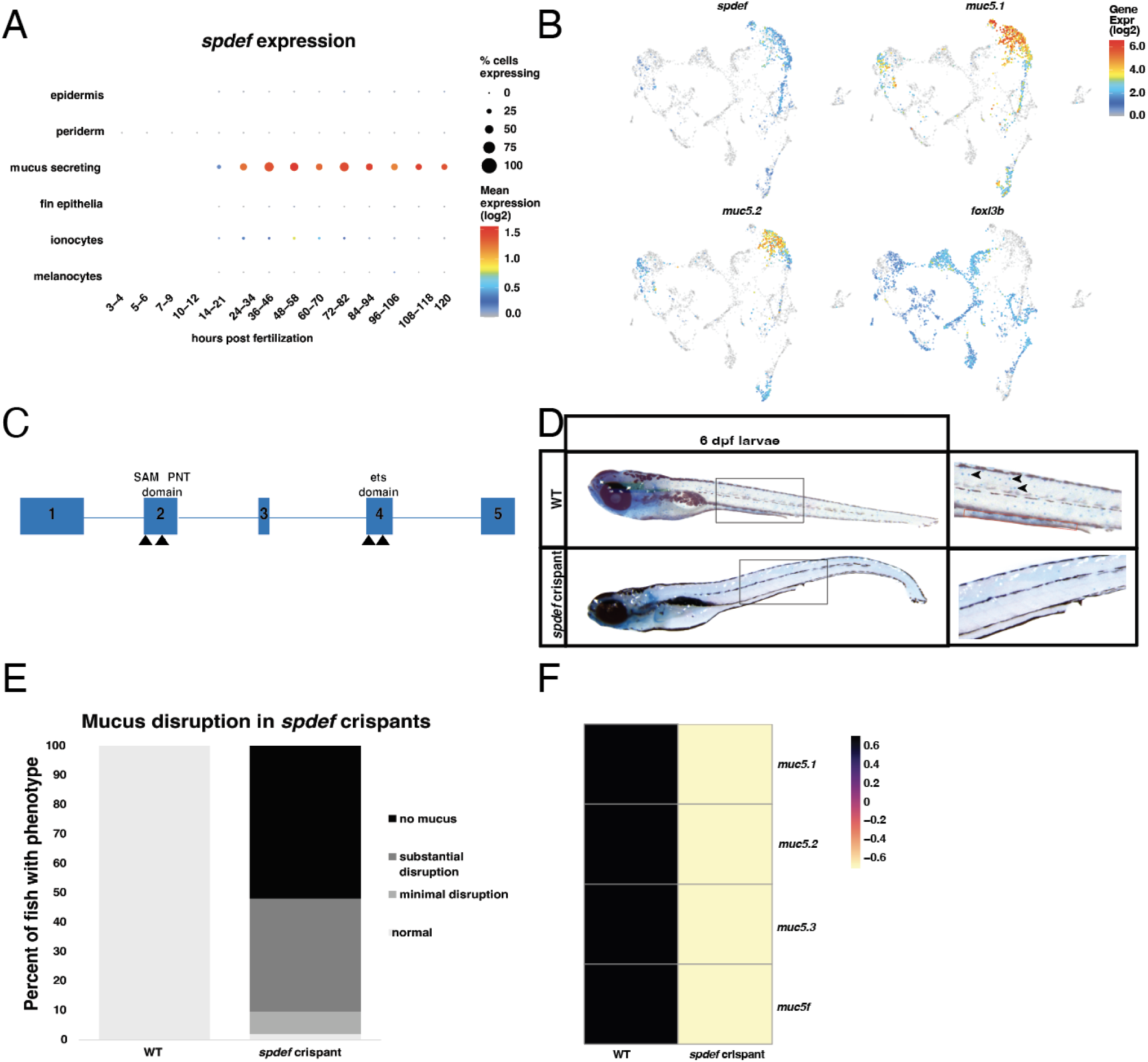
*spdef* crispants lack mucus. (A) Expression of *spdef* occurs in mucus-secreting cells of the epidermis beginning around 14 hpf. Made with DaniocellDesktop (Evans and Farrell 2025) (B) UMAP of mucus-secreting cells showing expression of *spdef* and the skin mucin genes *muc5*.*1* and *muc5*.*2*, as well as the ionocyte specific gene *foxi3b*. Made with DaniocellDesktop (Evans and Farrell 2025). (C) Schematic showing targeting of CRISPR gRNAs to the spdef gene. Two gRNAs target the functional SAM PNT domain in exon 2 and two additional gRNAs target the functional ets domain in exon 4. (D) Representative image of Alcian blue staining in 6 dpf larval zebrafish in Cas9-injected (labeled WT) and *spdef* crispants. Blue speckles (indicated with arrowheads) are mucocytes, intestinal mucocytes are highlighted by a red box, and all other mucocytes are epidermal mucocytes. (E) Mucus depletion categorization for 6 dpf Cas9-only injected (WT) and *spdef* crispant larvae. Categories are no mucus (no blue speckles), substantial disruption (1-10 speckles), minimal disruption (10-50 speckles), no disruption (>50 speckles) n = 81, 71, C = 4. (F) Decreased skin mucin gene expression in *spdef* crispants compared to Cas9-only injected (WT). Measured from the indicated genotypes using RNAseq on 6 dpf embryos after mock exposure. RNA was collected 5 h post mock UVB exposure. Gene expression is scaled by rows.

## REFERENCES

Aniseia, Yttria, Eric Waltari, Hejin Huang, et al. 2026. “RNAquarium: An Archive-Scale Atlas of Zebrafish Gene Expression Coupled with Pan-Taxonomic Profiling Reveals Diverse Viral Drivers of Transcriptomic States.” In bioRxiv. BioRxiv, June 19. 10.64898/2026.06.17.732950.

Behrendt, Lars, Maria E. Jönsson, Jared V. Goldstone, and John J. Stegeman. 2010. “Induction of Cytochrome P450 1 Genes and Stress Response Genes in Developing Zebrafish Exposed to Ultraviolet Radiation.” Aquatic Toxicology (Amsterdam, Netherlands) 98 (1): 74–82.

Blighe, K., S. Rana, and M. Lewis. 2019. “EnhancedVolcano: Publication-Ready Volcano Plots with Enhanced Colouring and Labeling.” R Package Version. https://scholar.google.com/citations?user=OSyxVocAAAAJ&hl=en&oi=sra.

Braun, C., R. Reef, and U. E. Siebeck. 2016. “Ultraviolet Absorbing Compounds Provide a Rapid Response Mechanism for UV Protection in Some Reef Fish.” Journal of Photochemistry and Photobiology. B, Biology 160 (July): 400–407.

Brenner, Michaela, and Vincent J. Hearing. 2008. “The Protective Role of Melanin against UV Damage in Human Skin.” Photochemistry and Photobiology 84 (3): 539–549.

Chatterjee, Nimrat, and Graham C. Walker. 2017. “Mechanisms of DNA Damage, Repair, and Mutagenesis: DNA Damage and Repair.” Environmental and Molecular Mutagenesis 58 (5): 235–263.

Chen, Gang, Thomas R. Korfhagen, Yan Xu, et al. 2009. “SPDEF Is Required for Mouse Pulmonary Goblet Cell Differentiation and Regulates a Network of Genes Associated with Mucus Production.” The Journal of Clinical Investigation 119 (10): 2914–2924.

Chuang, Ling-Shiang, Joshua Morrison, Nai-Yun Hsu, et al. 2019. “Zebrafish Modeling of Intestinal Injury, Bacterial Exposures and Medications Defines Epithelial in Vivo Responses Relevant to Human Inflammatory Bowel Disease.” Disease Models & Mechanisms 12 (8): dmm037432.

Cockell, C. S., and J. Knowland. 1999. “Ultraviolet Radiation Screening Compounds.” Biological Reviews of the Cambridge Philosophical Society 74 (3): 311–345.

Concordet, Jean-Paul, and Maximilian Haeussler. 2018. “CRISPOR: Intuitive Guide Selection for CRISPR/Cas9 Genome Editing Experiments and Screens.” Nucleic Acids Research 46 (W1): W242– W245.

Dunlap, Walter C., and J. Malcolm Shick. 1998. “Review—ultraviolet Radiation□absorbing Mycosporine□like Amino Acids in Coral Reef Organisms: A Biochemical and Environmental Perspective.” Journal of Phycology 34 (3): 418–430.

Evans, Alicia, and Jeffrey A. Farrell. 2025. “DaniocellDesktop: For Interactive Reanalysis of Wild-Type Zebrafish Single-Cell Genomic Data.” Zebrafish 22 (6): 230–232.

Gao, Qunjie, and Ferran Garcia-Pichel. 2011. “Microbial Ultraviolet Sunscreens.” Nature Reviews. Microbiology 9 (11): 791–802.

Görünmek, Mihraç, Bariş Ballik, Zeynep Elibol Çakmak, and Turgay Çakmak. 2024. “Mycosporine-like Amino Acids in Microalgae and Cyanobacteria: Biosynthesis, Diversity, and Applications in Biotechnology.” Algal Research 80 (103507): 103507.

Hawkes, J. W. 1974. “The Structure of Fish Skin. I. General Organization.” Cell and Tissue Research 149 (2): 147–158.

Jevtov, Irena, Tore Samuelsson, Grace Yao, Adam Amsterdam, and Katharina Ribbeck. 2014. “Zebrafish as a Model to Study Live Mucus Physiology.” Scientific Reports 4 (1): 6653.

Kapp, Friedrich G., Julie R. Perlin, Elliott J. Hagedorn, et al. 2018. “Protection from UV Light Is an Evolutionarily Conserved Feature of the Haematopoietic Niche.” Nature 558 (7710): 445–448.

Karentz, Deneb, and Louise H. Lutze. 1990. “Evaluation Ofbiologically Harmful Ultraviolet Radiation in Antarctica with a Biological Dosimeter Designed for Aquatic Environments.” Limnology and Oceanography 35 (3): 549–561.

Keating, Margaret K., Julio Jaime, Abigail Mumme-Monheit, et al. 2026. “Early Ultraviolet Exposure Disrupts Late Dorsal Fin Skeletal Development in Zebrafish.” Developmental Dynamics, no. dvdy.70133 (April). 10.1002/dvdy.70133.

Lange, Merlin, Alejandro Granados, Shruthi VijayKumar, et al. 2024. “A Multimodal Zebrafish Developmental Atlas Reveals the State-Transition Dynamics of Late-Vertebrate Pluripotent Axial Progenitors.” Cell 187 (23): 6742–6759.e17.

Lu, Yu-Fen, Da-Wei Liu, I. Chen Li, et al. 2021. “Delta/Jagged-Mediated Notch Signaling Induces the Differentiation of agr2-Positive Epidermal Mucous Cells in Zebrafish Embryos.” PLoS Genetics 17 (12): e1009969.

McNamara, M. E., V. Rossi, T. S. Slater, et al. 2021. “Decoding the Evolution of Melanin in Vertebrates.” Trends in Ecology & Evolution 36 (5): 430–443.

Mueller, Kaspar P., and Stephan C. F. Neuhauss. 2014. “Sunscreen for Fish: Co-Option of UV Light Protection for Camouflage.” PloS One 9 (1): e87372.

Osborn, Andrew R., Khaled H. Almabruk, Garrett Holzwarth, et al. 2015. “Abstract.” Preprint, ELife Sciences Organisation, Ltd. 10.7554/elife.05919.001.

Plack, P. A., N. W. Fraser, P. T. Grant, C. Middleton, A. I. Mitchell, and R. H. Thomson. 1981. “Gadusol, an Enolic Derivative of Cyclohexane-1,3-Dione Present in the Roes of Cod and Other Marine Fish. Isolation, Properties and Occurrence Compared with Ascorbic Acid.” The Biochemical Journal 199 (3): 741–747.

Rajavelu, Priya, Gang Chen, Yan Xu, Joseph A. Kitzmiller, Thomas R. Korfhagen, and Jeffrey A. Whitsett. 2015. “Airway Epithelial SPDEF Integrates Goblet Cell Differentiation and Pulmonary Th2 Inflammation.” The Journal of Clinical Investigation 125 (5): 2021–2031.

Rice, Marlen C., Jordan H. Little, Dale L. Forrister, Julane Machado, Nathan L. Clark, and James A. Gagnon. 2023. “Gadusol Is a Maternally Provided Sunscreen That Protects Fish Embryos from DNA Damage.” Current Biology 33 (15): 3229–3237.e4.

Schindelin, Johannes, Ignacio Arganda-Carreras, Erwin Frise, et al. 2012. “Fiji: An Open-Source Platform for Biological-Image Analysis.” Nature Methods 9 (7): 676–682.

Stierner, U., I. Rosdahl, A. Augustsson, and B. Kågedal. 1989. “UVB Irradiation Induces Melanocyte Increase in Both Exposed and Shielded Human Skin.” The Journal of Investigative Dermatology 92 (4): 561–564.

Sur, Abhinav, Yiqun Wang, Paulina Capar, Gennady Margolin, Morgan Kathleen Prochaska, and Jeffrey A. Farrell. 2023. “Single-Cell Analysis of Shared Signatures and Transcriptional Diversity during Zebrafish Development.” Developmental Cell 58 (24): 3028–3047.e12.

Takasugi, Paige R., Shengzhou Wang, Kimberly T. Truong, et al. 2022. “Orthogonal CRISPR-Cas Tools for Genome Editing, Inhibition, and CRISPR Recording in Zebrafish Embryos.” Genetics 220 (1): iyab196.

Teai, Taivini, Jacqui H. Drollet, Jean-Pierre Bianchini, Aimé Cambon and Paul M. V. Martin. 1998. “Occurrence of Ultraviolet Radiation-Absorbing Mycosporine-like Amino Acids in Coral Mucus and Whole Corals of French Polynesia.” Marine & Freshwater Research 49 (2): 127–132.

Van Der Leun, Jan C. 2004. “The Ozone Layer.” Photodermatology, Photoimmunology & Photomedicine 20 (4): 159–162.

Wickham, Hadley, Mara Averick, Jennifer Bryan, et al. 2019. “Welcome to the Tidyverse.” Journal of Open Source Software 4 (43): 1686.

Zamzow, Jill P., Ulrike E. Siebeck, Maxi J. Eckes, and Alexandra S. Grutter. 2013. “Ultraviolet-B Wavelengths Regulate Changes in UV Absorption of Cleaner Fish Labroides Dimidiatus Mucus.” PloS One 8 (10): e78527.

